# PITX1 protein interacts with ZCCHC10 to regulate *hTERT* mRNA transcription

**DOI:** 10.1101/640508

**Authors:** Takahito Ohira, Hirotada Kojima, Yuko Kuroda, Daigo Inaoka, Mitsuhiko Osaki, Hideki Wanibuchi, Futoshi Okada, Mitsuo Oshimura, Hiroyuki Kugoh

## Abstract

Telomerase is a ribonucleoprotein ribonucleic enzyme that is essential for cellular immortalization *via* elongation of telomere repeat sequences at the end of chromosomes. Human telomerase reverse transcriptase (*hTERT*), the catalytic subunit of telomerase holoenzyme, is a key regulator of telomerase activity. Telomerase activity, which has been detected in the majority of cancer cells, is accompanied by *hTERT* expression, resulting in continuous cell division that is a crucial factor for malignant transformation. Thus, *hTERT* is an attractive target for cancer-specific treatments. We previously reported that pared-like homeodomain 1 (*PITX1*) is a negative regulator of *hTERT* through direct binding to the *hTERT* promoter. However, the mechanism by which the function of *PITX1* contributes to transcriptional silencing of the *hTERT* gene remains to be clarified. Here, we show that PITX1 and zinc finger CCHC-type containing 10 (ZCCHC10) proteins cooperate to facilitate the transcriptional regulation of the *hTERT* gene by functional studies *via* FLAG pull-down assay. Co-expression of *PITX1* and *ZCCHC10* resulted in inhibition of *hTERT* transcription, in melanoma cell lines, whereas mutate-deletion of homeodomain in PITX1 that interact with ZCCHC10 did not induce similar phenotypes. In addition, ZCCHC10 expression levels showed marked decrease in the majority of melanoma cell lines and tissues. Taken together, these results suggest that ZCCHC10-PITX1 complex is a new functional unit that suppresses *hTERT* transcription, and plays a crucial role in cancer development through regulation of telomerase activity.

## Introduction

Telomerase ribonucleic enzyme is associated with cellular immortalization and contributes to elongation of telomere repeat sequences on the end of chromosomes, making it a key factor in cancer development [1,2]. Human telomerase consists of essential enzyme subunits; the protein catalytic subunit human telomerase reverse transcriptase (*hTERT*) and the RNA subunit, human telomerase RNA component (*hTERC*) and the accessory proteins dyskerin, NOP10, NHP2 and GAR1 [3,4]. The expression of *hTERT* is critical for telomerase enzyme activity. Indeed, ectopic *hTERT* expression in telomerase-negative normal cells can extend lifespan and establish immortalized cell lines *via* elongation of telomeres [5,6]. Expression of *hTERT* is down-regulated in most human adult somatic cells, except in germ cells and some stem cells. On the other hand, its expression was detected in the majority of human cancer cells (around 85-90%) [7,8]. This is consistent with telomerase conferring a strong selective advantage for continued growth of malignant cells, suggesting that telomerase activity is essential for most cancer cell immortalization and it may be possible to inhibit of cancer development by the control of *hTERT* expression. Although it is known that expression of *hTERT* is regulated by various activating and repressing transcription factors and epigenetic modification [9,10], the underlying molecular mechanisms that are involved in regulation of *hTERT* transcription during cellular differentiation and cancer development remains unclear.

We previously confirmed that human chromosomes 3, 5, and 10 carry *hTERT* regulatory genes using microcell mediated chromosome transfer (MMCT) [11]. In particular, we identified paired-like homeodomain 1 (*PITX1*) as a novel *hTERT* suppressor gene, located on human chromosome 5 by a combination of MMCT and gene expression profile analysis. *PITX1* regulates *hTERT* transcription through binding to its promoter [12,13]. *PITX1* was originally identified as a transcription factor gene that is able to activate pituitary transcription of a pro-opiomelanocortin gene. *PITX1*, which belongs to bicoid-related homeobox genes, plays a role in the development of the Rathke pouch and adult pituitary [14], and consists of an N-terminal homeodomain and a C-terminal Otp-Aristaless-Rax (OAR) domain. The homeodomain has DNA binding function and also acts as a protein-protein interaction for target molecule [15,16). On the other hand, the OAR domain may modulate transactivation capacities or be involved in protein-protein interactions [17,18]. *Pitx1* knockout mice developed fetuses with abnormal hindlimbs, thus suggesting that it regulates the developmental limb [19]. In addition, *PITX1* is known as a tumor suppressor gene that inhibits the *RAS* pathway through Ras protein activator-like 1 (*RASAL1*), which is a member of the Ras GTPase-activating protein family [20] and induces activation of *p53* transcription [21]. Furthermore, we provided important evidence that PITX1 directly binds to specific PITX1 response element sites in the *hTERT* promoter region, resulting in telomerase inhibition [13]. Downregulation of *PITX1* is observed in various cancers including malignant melanoma, oral, gastric, colon, lung, and bladder cancers [20,22–26]. Collectively, this evidence suggests that *PITX1* plays a crucial role in cancer development, though telomerase-dependent pathways. Interestingly, the introduction of an intact human chromosome 5 into melanoma A2058 cells more strongly suppressed *hTERT* transcription when compared with *PITX1* cDNA-overexpressing clones [12,27]. Therefore, human chromosome 5 carries one or more genes that are involved in the suppression of *hTERT* transcription, in addition to the *PITX1* gene.

The zinc fingers Lys-Cys-His-Cys-type 10 (*ZCCHC10*) gene belongs to the CCHC-type zinc finger nucleic acid binding protein family. DNA methylation level of CpG site in *ZCCHC10* gene using offspring cord blood DNA was showed potentially *ZCCHC10* related to apoptosis, tumorigenesis and inflammation pathways [28]. In addition, ZCCHC10 protein level is down-regulated in atopic dermatitis patients-derived serum [29]. However, the functional role of *ZCCHC10* gene in tumorigenesis is poorly understood. ZCCHC10 contains a single CCHC domain, which is known that mediate protein-protein interactions. For example, zinc finger protein FOG family member 1 (*FOG1*) contains five CCHC domain and each CCHC domain bind to GATA bunding protein 1 (*GATA1*) [30]. FOG1-GATA1 complexes function as activators for several genes, which required for normal erythroid differentiation and megakaryocyte maturation [31]. Taken together, these previously studies suggest that the CCHC domain in ZCCHC10 have the potential to interact with other biomolecules containing transcription factors and the complex may have a function of regulation for tumorigenesis related genes.

In this study, we identified ZCCHC10 as a protein that interacted with PITX1, and interaction sites were mapped to the CCHC domain of ZCCHC10 and the homeodomain of PITX1. Moreover, expression levels of *ZCCHC10* in human melanoma cells were significantly lower when compared with normal melanocyte cells. In addition, overexpression of both the *PITX1* and *ZCCHC10* genes showed significant suppression of *hTERT* transcription when compared to that of each gene in melanoma cell lines. Thus, our findings of the *PITX1* complex that regulates *hTERT* transcription may facilitate understanding of the molecular mechanisms involved in telomerase dependent cellular replicative senescence or cancer development throughout the *hTERT* signaling network.

## Results

### Identification of novel proteins that formed complex with PITX1 in *hTERT*-negative cells

Cellular immortalization of all human tumors requires maintenance of telomere length, which needs reactivation of telomerase enzyme (telomerase dependent pathways) or activation of telomerase independent Alternative Lengthening of Telomere (ALT) pathways. We propose that functional PITX1 complexes that regulate *hTERT* transcription are present in telomerase-negative tumor cells in which *hTERT* transcription is not detectable, but not in telomerase-positive tumor cells. To determine protein interactions with PITX1, we performed pull-down assay using *hTERT*-positive HeLa229, 293T cells [32,33] and *hTERT*-negative ALT U2OS [34] cells with FLAG-tagged PITX1. We used control cells that expressed only FLAG-tag protein (Fig 1A). First, FLAG-tagged PITX1 was transfected into HeLa229, 293T and U2OS cells, respectively. Enrichment of FLAG-tagged PITX1 protein in cell lysates obtained by immunoprecipitation with anti-FLAG antibody was confirmed using western blotting analysis (Fig 1B). A band at 49 kDa corresponding to FLAG-tagged PITX1 was detected in these cancer cells, but not in control cells, on Coomassie brilliant blue (CBB) and silver stained gels (Fig 1C). nanoLC/MS/MS-based proteomic analysis of FLAG immunoprecipitation extracts obtained from these cell lines yielded 27 (HeLa229), 34 (293T), and 101 (U2OS) proteins, respectively (Fig 1D and Supplementary table S1). Ultimately, 57 proteins were identified only in *hTERT*-negative U2OS cells (Fig 1D). Therefore, it is likely that PITX1-interacting proteins function as suppressors of *hTERT* transcription.

**Fig 1.**
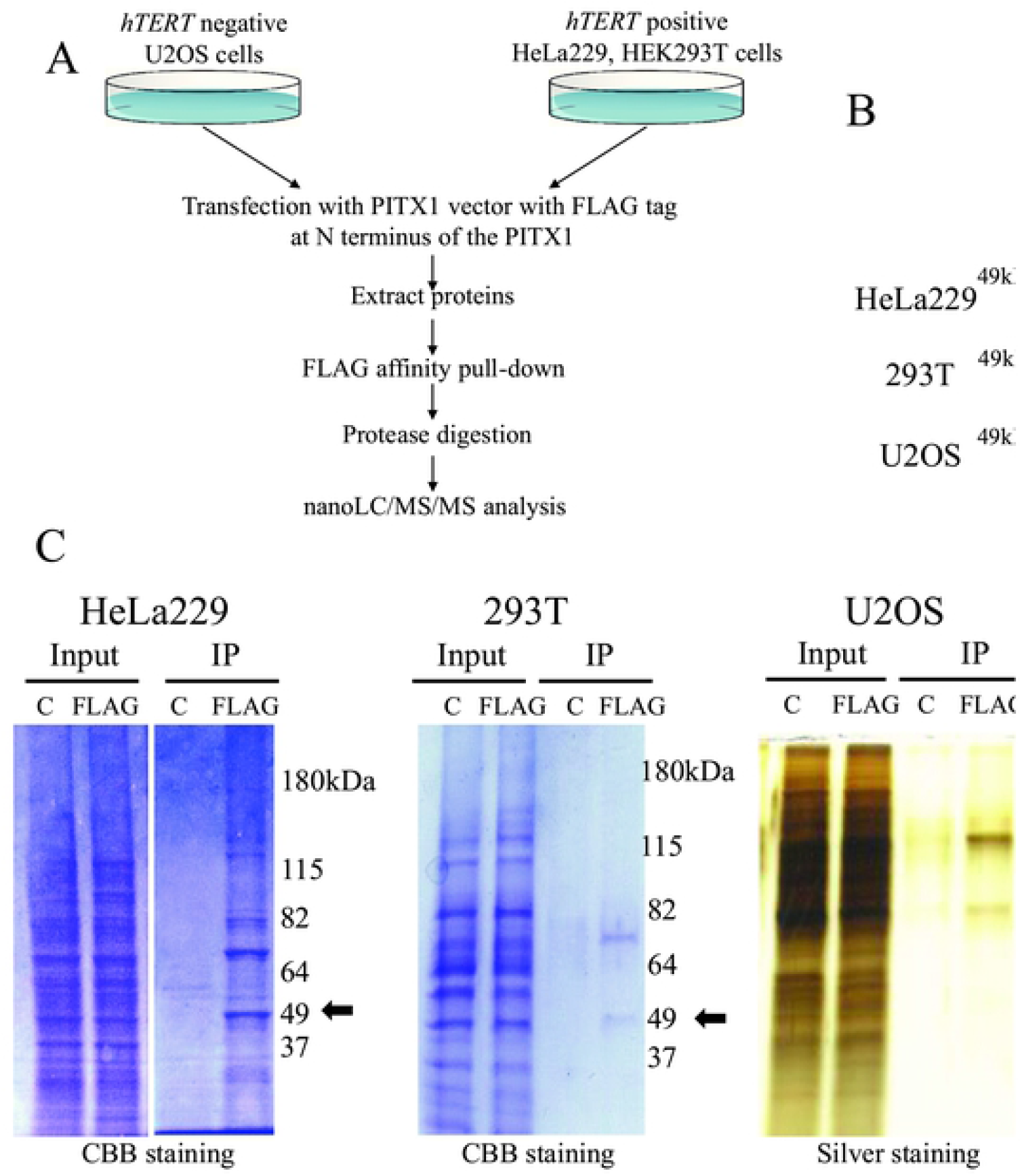
Overview of analysis of proteins associated with affinity purified FLAG-PITX1 by mass spectrometry. (A) Purification scheme and composition of PITX1 complex. PITX1-associated proteins were affinity purified from whole cell extracts of FLAG-PITX1-expressing *hTERT*-positive HeLa229 and HEK293T cells, and *hTERT*-negative U2OS cells using anti-FLAG M2 affinity gel. Bound proteins were eluted with 300 ng/μl FLAG peptide and treated with protease to digestion. Then, digestion proteins were preformed nanoLC/MS/MS assay to identify pull-down proteins. (B) Immunoprecipitation of FLAG-PITX1 with anti-FLAG was followed by western blotting using anti-PITX1 antibody. Left panel shows input samples for immunoprecipitated proteins. Right panel shows proteins immunoprecipitated by FLAG-antibody from FLAG-PITX1-overexpressing cells and control cells. (C) CBB (HeLa229 and 293T cells) and silver staining (U2OS) of the PITX1-associated protein after SDS-PAGE (10%) is shown. FLAG empty vector-expressing cells using the same protocol (negative control) are shown. Black arrow indicates FLAG-PITX1 protein at around 49 kDa. C: empty vector control lysate. FLAG: FLAG-PITX1 expression lysate. (D) Immunoaffinity profiling of *hTERT* negative expressing U2OS cells, as compared with *hTERT* positive expressing HeLa229 and 293T cells by nanoLC/MS/MS-based proteomic approaches.

### Validation of PITX1-interacting proteins

We previously reported that microcell hybrid clones obtained by the introduction of human chromosome 5 completely suppressed *TERT* transcription [12]. On the other hand, the suppression effects of *hTERT* transcription in transfectant cells that were overexpressed by introduction of *PITX1* cDNA were weak when compared to those in microcell hybrid cells with human chromosome 5 in A2058 human melanoma cells [12,27]. This suggests that PITX1 interacts with other proteins on human chromosome 5 to form a functional complex essential for suppression of *hTERT* transcription. Therefore, we focused on genes located on chromosome 5 as novel PITX1 interacting proteins. As a result, among 57 candidate genes, six genes localized on human chromosome 5 were identified as PITX1-interacting proteins in *hTERT*-negative U2OS cells (Table 1 and Supplementary Fig S1). Interestingly, three of these genes, namely, heterogeneous nuclear ribonucleoprotein A0 (*hnRNP A0*), *ZCCHC10* and 75 kDa glucose-regulated protein (*HSPA9*), were localized in the peripheral *PITX1* region (5q31.1). We first aimed to identity PITX1-interacting proteins using the Human Protein Atlas (HPA) database, which includes information on gene expression in normal and cancer cells and tissues (http://www.proteinatlas.org/). Among these, *ZCCHC10* showed lower expression in melanoma tissue samples when compared with normal melanocyte cells. To further validate the authenticity of the PITX1-ZCCHC10 complex indicated by nanoLC/MS/MS analysis, we investigated whether ZCCHC10 is present in the PITX1 immunocomplex in FLAG-PITX1-expressing U2OS cells by western blotting analysis using anti-ZCCHC10 antibody. Western blotting detected bands suggesting that ZCCHC10 protein forms a complex with FLAG-PITX1 protein (Fig 2). Thus, ZCCHC10 directly or indirectly interacts with PITX1.

**Fig 2.**
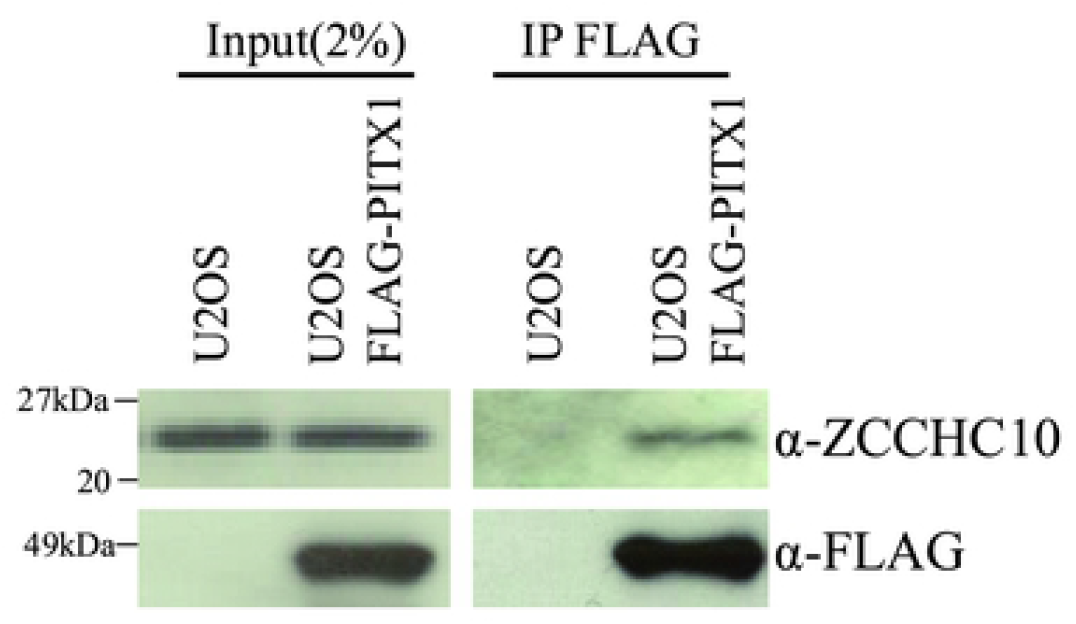
PITX1 associated with endogenous ZCCHC10 protein in U2OS cells. Confirmation of pull-down assay results by immunoprecipitation using FLAG-tagged PITX1 in *hTERT*-negative cell line. U2OS cells transfected FLAG-PITX1 vector were immunoprecipitated with an anti-FLAG antibody and immunoblotted with anti-FLAG and anti-ZCCHC10 antibody. Left panel shows input samples for immunoprecipitated proteins. Right panel shows proteins immunoprecipitated by FLAG-antibody from FLAG-PITX1-overexpressing U2OS cells and control cells.

**Table 1.**
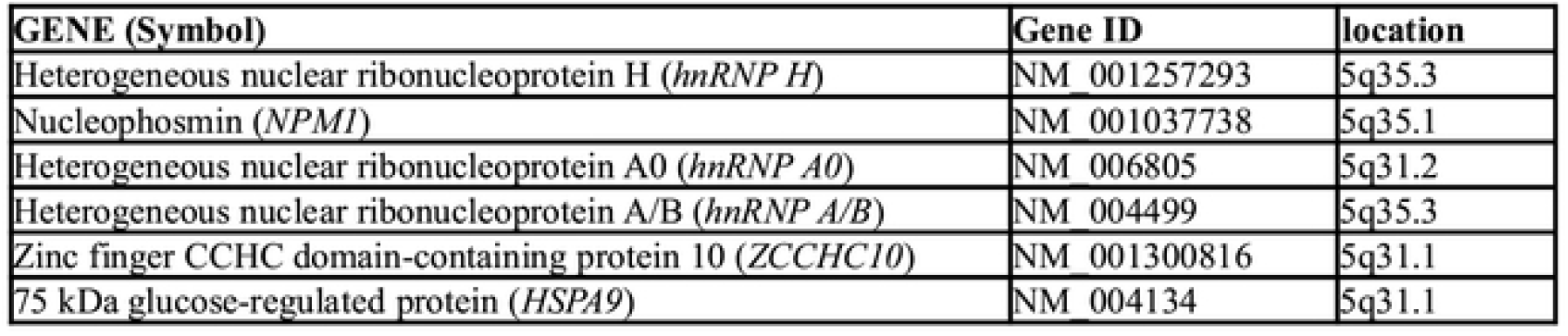
*PITX1*-associated genes in *TERT* negative U2OS cells located on human chromosome 5.

| **GENE (Symbol)** | **Gene ID** | **location** |
| --- | --- | --- |
| Heterogeneous nuclear ribonucleoprotein H (*hnRNP H*) | NM_001257293 | 5q35.3 |
| Nucleophosmin (*NPM1*) | NM_001037738 | 5q35.1 |
| Heterogeneous nuclear ribonucleoprotein A0 (*hnRNP A0*) | NM_006805 | 5q31.2 |
| Heterogeneous nuclear ribonucleoprotein A/B (*hnRNP A/B*) | NM_004499 | 5q35.3 |
| Zinc finger CCHC domain-containing protein 10 (*ZCCHC10*) | NM_001300816 | 5q31.1 |
| 75 kDa glucose-regulated protein (*HSPA9*) | NM_004134 | 5q31.1 |

### Expression profiles of ZCCHC10 expression level in melanoma cell lines and tissues

We next performed western blotting analysis to investigate the expression profiles of *ZCCHC10* in human melanoma cell lines. The expression levels of ZCCHC10 proteins were significantly reduced in all six melanoma cell lines (A2058, SKMEKL28, G361, HMV1, CRL1597 and GAK) when compared to normal human epidermal melanocyte cells (Fig 3A). Furthermore, qRT-PCR analysis indicated that *ZCCHC10* mRNA expression was markedly decreased in 11 out of 12 human clinical melanoma specimens when compared with normal melanocyte cells (Fig 3B). Moreover, our previous data demonstrated that *PITX1* expression is also down-regulated in melanomas [22]. These data suggest that both *PITX1* and *ZCCHC10* play a crucial role as tumor suppressor genes and the expression profiles of those genes are involved in melanoma development through dysregulation of *hTERT* transcription.

**Fig 3.**
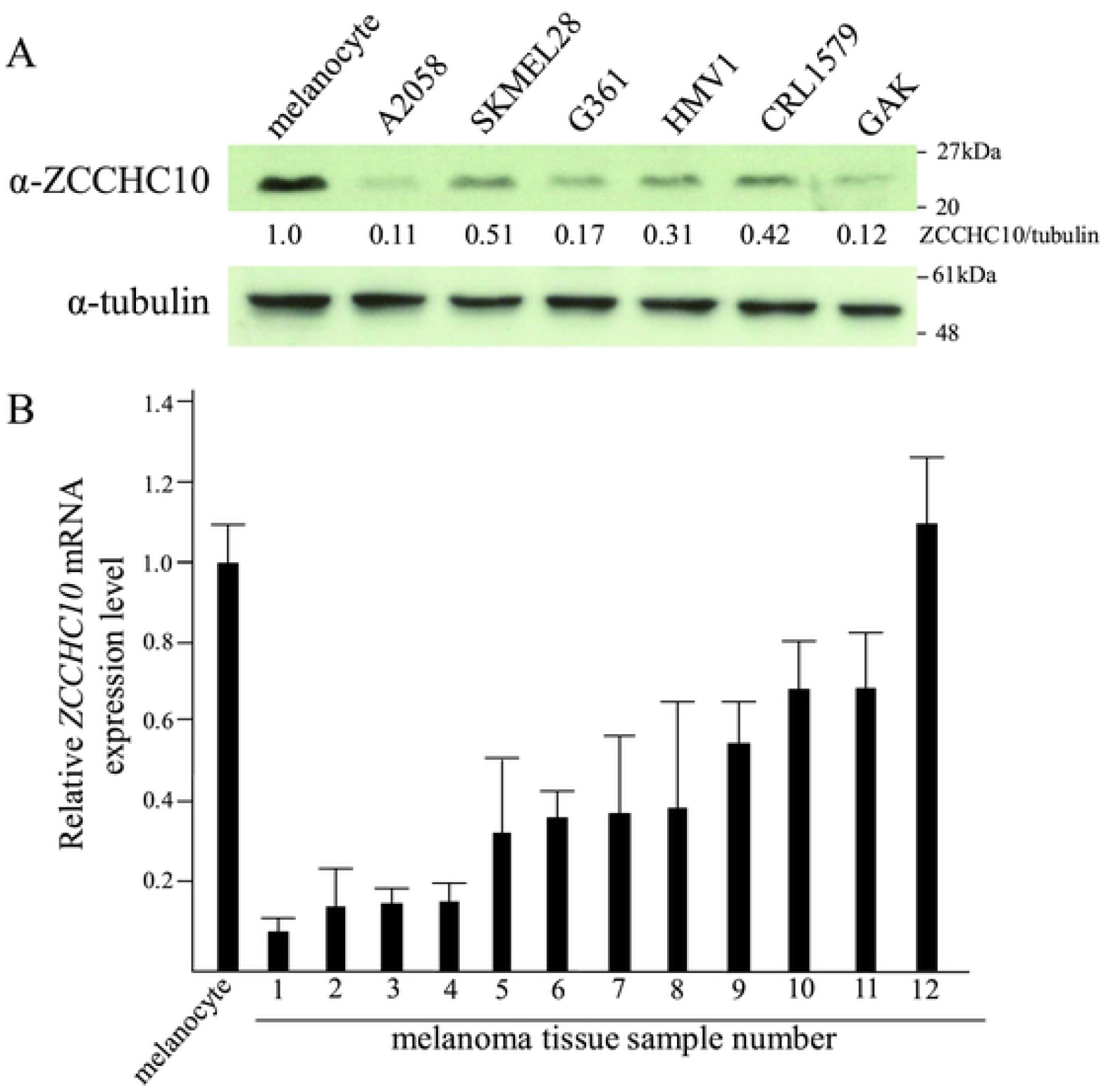
Downregulation of *ZCCHC10* in melanoma cells. Expression analysis of *ZCCHC10* in melanoma cell lines and tissue sample. (A) Expression levels of ZCCHC10 were analyzed by western blotting. A2058, SKMEL28, G361, HMV1, CRL1579, and GAK are melanoma cell lines. Melanocytes were used as normal cells. Expression levels of ZCCHC10 were normalized against levels of β-tubulin. (B) qRT-PCR analysis of *ZCCHC10* expression levels in human melanoma clinical tissue samples relative to those in melanocytes. Data were normalized against *GAPDH* mRNA. Expression in melanocytes was arbitrarily at set 1. Bars correspond to means ±S.D. of three independent experiments.

### ZCCHC10-PITX1 complex suppress *hTERT* expression level

In order to investigate the suppression effects of *hTERT* transcription by the PITX1-ZCCHC10 complex, we transiently co-transfected A2058 cells with expression vectors containing the *PITX1* cDNA and *ZCCHC10* cDNA sequences. Transient overexpression of *PITX1* or *ZCCHC10* in A2058 cells reduced *hTERT* transcription to 84% or 82% of control cells, respectively. On the other hand, transient co-expression of *PITX1* and *ZCCHC10* vector induced a significant decrease in transcription to 59% of that in control cells (Fig 4A). To further examine whether PITX1 and ZCCHC10 form protein complexes, we carried out immunoprecipitation of cell lysates from FLAG-PITX1 or ZCCHC10 overexpressing A2058 cells using anti-PITX1 and ZCCHC10 antibodies. Bands corresponding to PITX1 and ZCCHC10 were detected by western blotting in both pull-downs (Fig 4B and C). These results suggest that PITX1 is able to form a complex with ZCCHC10, and is functionally involved in the suppression of *hTERT* transcription in A2058 cells.

**Fig 4.**
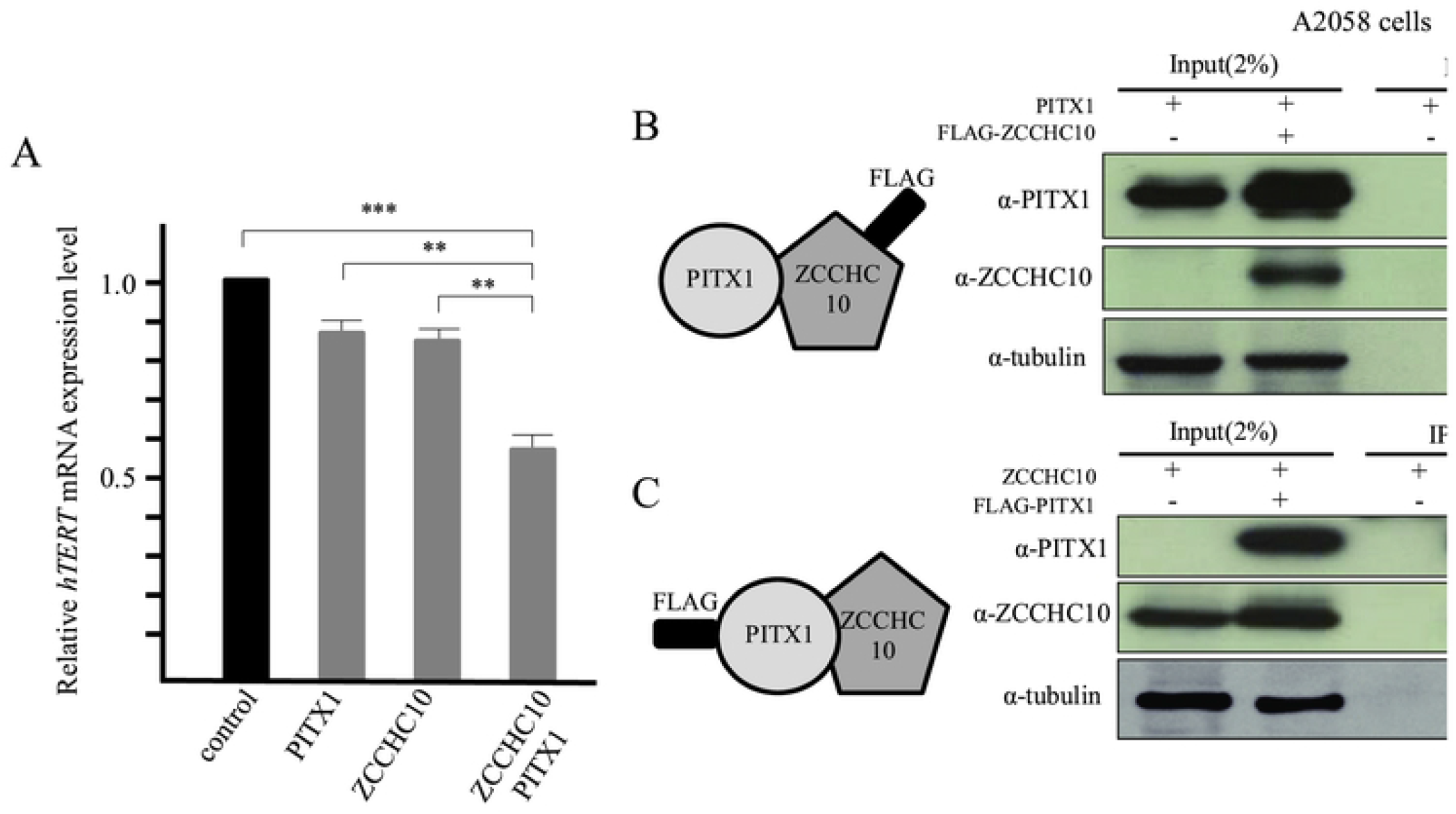
ZCCHC10/PITX1 complex suppressed *hTERT* mRNA expression in A2058 cells. (A) Co-transfection of *ZCCHC10* and *PITX1* in A2058 cells and qRT-PCR analysis of *hTERT* mRNA expression levels relative to empty vector-transfected control cells. Data were normalized against *GAPDH* mRNA. Expression level in control-vector cells was arbitrarily assigned as 1. Bars correspond to means ±S.D. of three independent experiments (***P<0.001, **P<0.01). (B) Total protein extracts from A2058 cells, transfected with FLAG-ZCCHC10 and PITX1 expression vectors, were subjected to immunoprecipitation with anti-FLAG M2 affinity gel followed by western blotting with anti-PITX1 and anti-ZCCHC10 antibodies. As a control, empty FLAG vector was transfected into A2058 cells. (C) Total protein extracts from A2058 cells, transfected with FLAG-PITX1 and ZCCHC10 expression vectors, were subjected to immunoprecipitation with anti-FLAG M2 affinity gel, followed by western blotting with anti-ZCCHC10 and anti-PITX1 antibodies. As a negative control, empty FLAG vector was transfected into A2058 cells.

### Homeodomain of PITX1 interacts with CCHC domain of ZCCHC10

To determine the protein-protein interaction sites between ZCCHC10 and a PITX1, we performed mutation analysis. PITX1 is a transcription factor that contains a homeodomain and an OAR, which are the predicted protein-protein interaction domain [15–17]. First, we prepared a FLAG-tagged PITX1 deleted homeodomain 1 (half of the N-terminal side of the homeodomain deleted: FLAG-PITX1 ΔHD1), FLAG-PITX1 homeodomain 2 (half of the C-terminal side of the homeodomain deleted: FLAG-PITX1 ΔHD2) and FLAG-PITX1 deleted OAR (OAR domain deleted: FLAG-PITX1 ΔOAR) protein expression plasmid vectors (Fig 5A). FLAG-tagged wild-type PITX1 (FLAG-PITX1 wt) or each FLAG-tagged mutation-type PITX1 vector was co-expressed with an expression vector containing the wild-type ZCCHC10 expression vector in A2058 cells. Immunoprecipitation analysis was performed using anti-FLAG antibody. Western blotting using anti-ZCCHC10 antibody showed co-precipitated ZCCHC10 with PITX1 wt and ΔOAR, but not with ΔHD1 and ΔHD2 (Fig 5B). This indicates that the homeodomain in PITX1 is essential for forming a protein complex with ZCCHC10.

**Fig 5.**
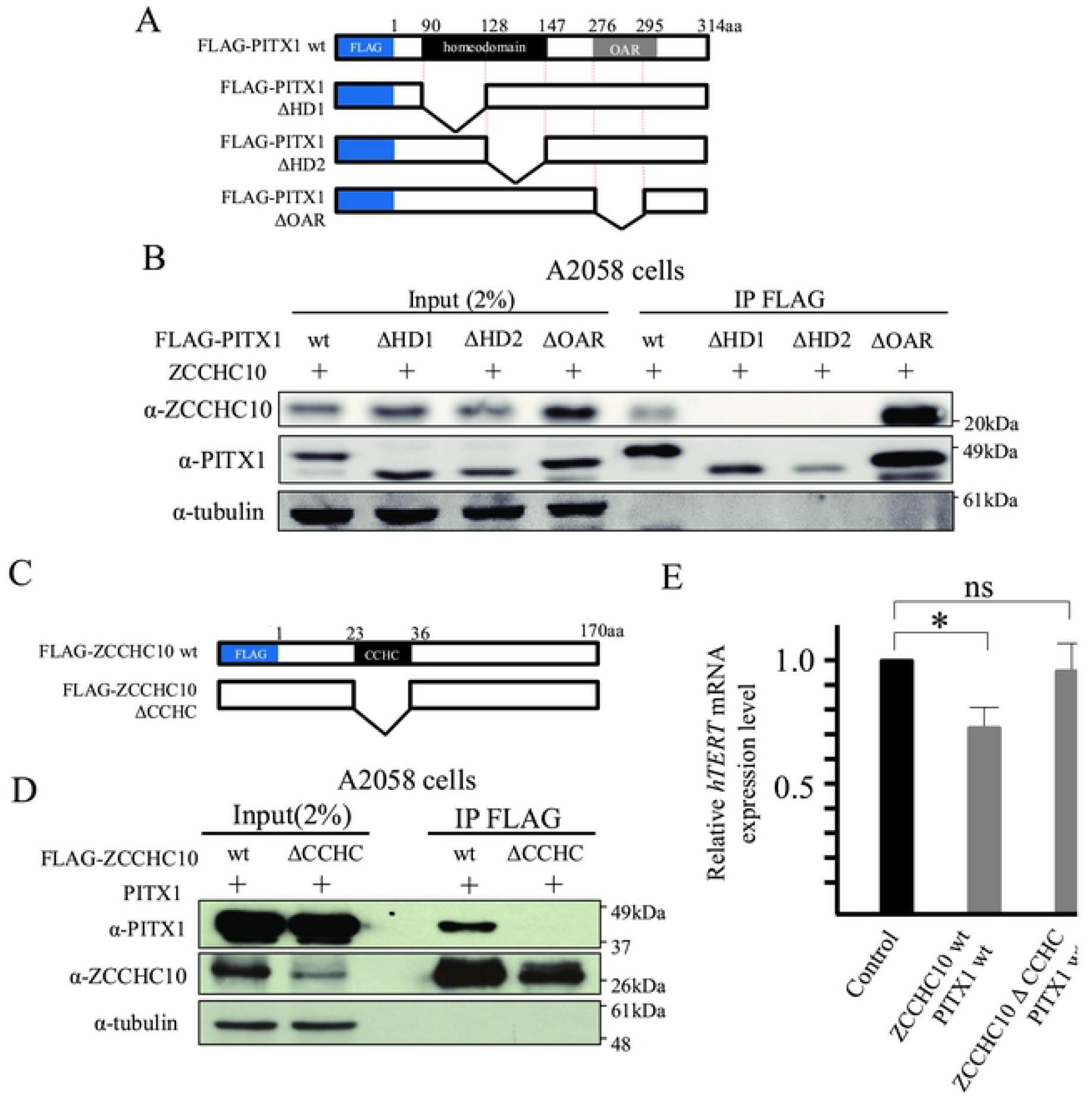
Homeobox domain of PITX1 binds the CCHC domain of ZCCHC10 *in vitro.* Identification of functional domain in PITX1 and ZCCHC10 protein, which is essential for the formation of protein complex for suppression of *hTERT* mRNA transcription. (A) PITX1 proteins used in this study with homeodomain and OAR domain. (B) Interactions between ZCCHC10 and PITX1 were examined by pull-down assays, using FLAG fusion proteins: FLAG-PITX1, FLAG-PITX1 ΔHD1, FLAG-PITX1 ΔHD2, FLAG-PITX1 ΔOAR. A2058 cells were transiently co-transfected with FLAG-PITX1 (or FLAG-PITX1 ΔHD1 or FLAG-PITX1 ΔHD2 or FLAG-PITX1 ΔOAR) and ZCCHC10 expression vector. Binding of ZCCHC10 to PITX1 was analyzed by western blotting using an anti-ZCCHC10 antibody. HD is the region of PITX1 that interacts with ZCCHC10. The OAR domain is not required for PITX1 interaction with ZCCHC10. (C) ZCCHC10 proteins used in this study with the CCHC domain. (D) Interactions between PITX1 and ZCCHC10 were examined by pull-down assay using FLAG fusion proteins: FLAG-ZCCHC10, FLAG-ZCCHC10 ΔCCHC. A2058 cells were transiently transfected with FLAG-ZCCHC10 (or FLAG-ZCCHC10 ΔCCHC) and PITX1 expression vector. Binding of PITX1 to ZCCHC10 was analyzed by western blotting using an anti-PITX1 antibody. The CCHC domain is the region of ZCCHC10 that interacts with PITX1. (E) Co-transfection of FLAG-ZCCHC10 ΔCCHC and PITX1 in A2058 cells and qRT-PCR analysis of *hTERT* mRNA expression levels relative to empty vector-transfected control cells. Data were normalized against *GAPDH* mRNA. The expression level in control-vector cells was arbitrarily assigned as 1. Bars correspond to means ±S.D. of three independent experiments (*P<0.05, ns: not significant).

On the other hand, the ZCCHC10 protein possesses the CCHC domain, which is known as a protein interaction domain. To determine the region of ZCCHC10 that interacted with PITX1, we performed a deletion analysis of the ZCCHC10 protein. We generated a FLAG-tagged ZCCHC10 plasmid vector, in which the CCHC domain was deleted (FLAG-ZCCHC10 ΔCCHC) (Fig 5C). FLAG-tagged wild-type ZCCHC10 (FLAG-ZCCHC10 wt) or FLAG-ZCCHC10 ΔCCHC was co-expressed with the expression vector coding the wild-type of PITX1 expression vector in A2058 cells. Immunoprecipitation analysis was performed using anti-FLAG antibody. Western blotting using anti-PITX1 antibody showed co-precipitation of PITX1 with wild-type ZCCHC10 protein, but not with ZCCHC10 ΔCCHC. This indicates that PITX1 forms a protein complex through the CCHC domain in ZCCHC10 (Fig 5D). Furthermore, we performed qRT-PCR analysis to investigate the effects of PITX1 and ZCCHC10 protein interaction regions on *hTERT* transcription. Expression levels of *hTERT* mRNA in A2058 cells co-transfected with ZCCHC10 wt and PITX1 wt expressing vectors were markedly decreased when compared to empty vector-transfected control cells (reduced *hTERT* transcription to 71% of control cells with empty vector). On the other hand, co-transfection to ZCCHC ΔCCHC with PITX1 wt vector in A2058 cells had no effect on *hTERT* transcription when compared to that in control cells (Fig 5E). These results suggest that interaction between PITX1 and ZCCHC10 proteins is mediated by CCHC on ZCCHC10 and that the homeodomain on PITX1 is essential for formation of a functional complex that regulates *hTERT* transcription.

## Discussion

*TERT*, which plays a crucial role in the regulation of telomerase activity, contributing to stem cell self-renewal and immortalization of malignant cells, is regulated by many different genes in response to a wide variety of oncogenic and suppressive signaling pathways. We previously identified *PITX1* as one of the *hTERT* suppressor, which directly binds to its promoter region [13]. In this study, we found that *ZCCHC10* is one of the components in *PITX1* complexes that significantly regulate *hTERT* transcription. *ZCCHC10* is also located in the same human chromosome region 5q31.1 as the *PITX1* gene. Recently, another group reported that a long noncoding RNA esophagus epithelial intergenic specific transcript (*Epist*) can down-regulate *hTERT* mRNA transcription [35]. Interestingly, the *Epist* RNA transcript is located 4.5-kbp downstream of the *PITX1* coding region. Furthermore, it has been reported that chromosome rearrangements that contain interstitial deletions and chromosome breakpoints were present on the long arm of chromosome 5 in acute myeloid leukemia (AML) and myelodysplasia (MDS) [36]. In particular, the most commonly deleted segment of 5q in MDS is within the 5q31 region [37]. This circumstantial evidence suggests that the human chromosome 5q31 region, which contains a negative regulator of *hTERT*, plays a significant role as the *hTERT* regulatory center.

In this study, three genes (*ZCCHC10, hnRNPA0*, and *HSPA9*), which were localized in the human chromosome 5q31.1-q31.2 region near *PITX1*, were identified as candidate proteins that interacted with *PITX1* using pull-down assay. *hnRNP A0* belongs to the A/B subfamily of ubiquitously expressed heterogeneous nuclear ribonucleoproteins (*hnRNPs*). The *hnRNAPs* are complexes of RNA and protein that localize in the cell nucleus during gene transcription and subsequent post-transcriptional modification of newly synthesized RNA (pre-mRNA). The *hnRNAPs* have important roles in multiple aspects of nucleic acid metabolism and in the regulation of different cellular processes, and some of them are reportedly associated with cancer development [38]. As little is known about the role of *hnRNP A0* in the maintenance of telomerase activity, including the regulation of *hTERT* transcription, functional analysis of this gene may provide future insights into the regulation telomerase. Other groups have previously reported that HSPA9 directly interacts with p53 protein and inhibits nuclear translocation and activation of *p53* mRNA transcription. In addition, targeting HSPA9-p53 complex with shRNA may induce p53-mediated apoptosis in hepatocellular carcinoma [39]. Therefore, *HSPA9* was identified as an oncogene through the physical inhibition of *p53.* On the other hand, we have provided evidence to suggest that HSPA9 protein interacts with PITX1 protein in *hTERT*-negative U2OS cells, which are known to express wild-type *p53* in response to radiation [40]. Thus, these results might suggest that the mechanism by which *PITX1* suppresses *HSPA9* oncogenic function involves the inhibition of telomerase activity. Interestingly, *PITX1* was identified as an activator of *p53* mRNA transcription *via* direct binding to the promoter [21]. It is possible that *PITX1* not only plays a role in regulating transcription *via* directly binding to the target gene promoter region, but may also play a role as a *p53* transcription factor and an inhibitor of *p53* negative regulator, ultimately controlling telomerase activity.

Homeodomain is one of the most important eukaryotic DNA-binding motifs and is highly conserved in sequence, structure, and mechanism of DNA binding. The homeodomain proteins, which include Nanog homeodomain protein (NANOG) and POU class 5 homeobox 1 (POU5F1 also known as OCT4), play an essential role in the development of pluripotent cells in the embryo, and in the self-renewal of embryonic stem (ES) cells and induced pluripotent stem (iPS) cells. These proteins are the transcription factors that regulate target genes, including epigenetic regulators directly binding to these promoter regions using homeodomain DNA-binding motifs [41]. However, little is known regarding the mechanisms by which these domains act in protein-protein interactions for target molecules. It has been reported that the interferon α (*INF-α*) promoter is regulated by *PITX1* in a manner similar to that of the *TERT* promoter. Interestingly, the homeodomains in PITX1 protein interacts with interferon regulatory transcription factor 3 (IRF3) and IRF7 proteins that control *INF-α* gene expression [15]. In addition, another group has shown that *PITX* families containing *PITX1, PITX2* and *PITX3* proteins interact *via* homeodomains with the DNA-binding domain of glial cell missing a (GCMa) proteins, which are known be transcription factors. The PITX-GCMa protein complex binds to the target promoter and influences GCMa-dependent promoter activity in a cell-specific manner [16]. Here, we found that interaction in ZCCHC10 protein through homeodomain in PITX1 controls *hTERT* mRNA transcription, suggesting that the homeodomain also plays a crucial role in regulating the target molecule. These data suggest that the homeodomain of *PITX* gene plays a distinctive role in transcriptional regulation of target genes when compared with other homeobox genes.

We reported here that *PITX1* interacted with *ZCCHC10*, resulting in significant suppression of *hTERT* transcription when compared to overexpression of *PITX1* or *ZCCHC10* alone. *PITX1* has been shown to activate the *p53* tumor suppressor gene, which regulates cell cycle progression, DNA integrity and cell survival, by directly binding to the *p53* promoter [21]. In addition, we previously showed that PITX1 binds to the *hTERT* promoter to suppress *hTERT* transcription [13]. Interestingly, *p53* is predicted to interact with ZCCHC10 based on high-throughput Protein-Protein Interaction Network data (http://genomenetwork.nig.ac.jp/). Furthermore, *p53* can also suppress *hTERT* transcription in various tumors, such as lung, prostate, and breast cancer cells [42]. These findings provide evidence that *ZCCHC10* plays an important role in the regulation of *hTERT* through *PITX1*- and p53-dependent pathways.

We previously showed that introduction of an intact human chromosome 5 markedly suppresses *hTERT* transcription in melanoma A2058 cells [12]. On the other hand, in this study, transient co-expression of *PITX1* and *ZCCHC10* induced a partial decrease in *hTERT* transcription (Fig 4A). It is possible that there are other negative regulatory factors and component proteins in the PITX1-ZCCHC10 complex on human chromosome 5 for functional *hTERT* control, which would ultimately lead to a large discrepancy in suppression of *hTERT* transcription in cancer cells. In fact, although *PITX1* is localized in the 5q31 region, we have previously found that a putative telomerase repressor gene was mapped to the 5p11 to 5p13 region by combination of functional analysis using chromosome engineering technology [43]. In this study, we found that at least six distinct PITX1-interacting proteins are present on human chromosome 5. Therefore, further studies that include identification of novel *hTERT* suppressor genes and functional analysis of protein or genes, should facilitate our understanding of the molecular mechanisms involved in the *hTERT* regulatory network *via PITX1*.

## Experimental procedures

### Cell culture

293T, A2058 and GAK cells were obtained from Japanese Collection of Research Bioresources Cell Bank (Osaka, Japan). SKMEL28, G361, HMV1 and CRL1579 cells were obtained from the Cell Resource Center for Biomedical Research, Institute of Development, Aging and Cancer, Tohoku University, Japan. HeLa229 and U2OS cells were obtained from American Type Culture Collection. Cells were cultured in Dulbecco’s Eagle’s medium (DMEM; Sigma, St. Louis, MO, USA) supplemented with 10% fetal bovine serum (HyClone, Logan, UT, USA). Human Epidermal Melanocyte cells (Invitrogen) were cultured in Medium 254 (Invirogen, Portland, OR, USA) supplemented with Human Melanocyte Growth supplement (Invitrogen). All cells were cultured at 37°C in a humidified incubator under 5% CO_2_. FLAG-PITX1 stable expression clones were maintained in DMEM supplemented with 10% FBS and G418 (HeLa229 and U2OS 500 μg/ml, 293T 600 μg/ml; Calbiochem, La Jolla, CA, USA). All Cell lines detected as Mycoplasma-free by MycoAlert mycoplasma detection kit (Lonza, Walkersville, MD, USA) and no more than 20 passage from the validated stocks.

### Plasmid vector

FLAG-tagged *PITX1* expression plasmids were as described previously [27]. pFLAG-ZCCHC10 was constructed by amplification of *ZCCHC10* cDNA from IMR90 cDNA by PCR using KOD plus DNA polymerase (TOYOBO, Osaka, Japan) and the following primer sequences; forward primer: 5’- GAAGATCTTATGGCGACTCCCATGCATCGGCTAA, reverse primer: 5’- GGGGTACCCTATTTCTTTTTCTTCTTCTTTGGT. Sequences were inserted into the Bgl Ⅱ/Acc65I (TOYOBO) digested FLAG-tagged vector pCMV-FLAG4 (Sigma). pCMV-FLAG4 was used as a control.

Non-tagged PITX1 or ZCCHC10 expression vector was inserted into the pCMV6-XL5 vector (Origene, Rockville, MD, USA). PITX1 ΔHD1, ΔHD2, ΔOAR and ZCCHC10 ΔCCHC expression plasmids were established using a mutagenesis kit (TOYOBO), according to the manufacturer’s instructions. The following primer sequences were used; PITX1 ΔHD1 forward primer: 5’-CGCGAGCGTAACCAGCAGCTGGAC, reverse primer: 5’-GCTCATGTCGGGGTAGCGGTTCCT, PITX1 ΔHD2 forward primer: 5’- ATGAGGGAGGAGATCGCCGTGTGG, reverse primer: 5’- CTGCTTCTTCTTCTTGGCTGGGTC, PITX1 ΔOAR forward primer: 5’- TTTGGCTACGGCGGCCTGCAGGGC, reverse primer: 5’- GTAGACGCTGTAGGGCGAGGCGGG, ZCCHC10 ΔCCHC forward primer: 5’- ACAGGAAAAAGAAAATACCTACAT, reverse primer: 5’- TCTTACATGTTGCTT ATTTGCTTC.

### Cell lysis and immunoprecipitation

Cells were harvested and resuspended in 5 ml of PBS at a concentration of 2×10^7^ cells/ml. HeLa229, 293T and U2OS cells stably expressing FLAG-tagged proteins were lysed in 5 ml of ice-cold lysis buffer [100 mM Tris HCl (pH 7.5), with 300 mM NaCl, 2 mM EDTA, 2% Triton X-100, and 400 μl of 25×EDTA-free protease inhibitors (Sigma) per 5 ml]. Soluble fractions from cell lysates were isolated by centrifugation at 15,000 × g for 30 minutes in a microfuge. For immunoprecipitation, 50 μl of anti-FLAG affinity gel (Sigma) was added to each lysate, followed by incubation with rotation for 24 hours at 4°C. Immunoprecipitates were washed three times with lysis buffer, and 50 μl of 150 μg/ml FLAG peptide was used to elute FLAG tagged proteins. Elution was performed at 4°C and lasted for 30 minutes for each elution. Gels were centrifuged for 1 min at 5,000 × g, and supernatants were transferred to fresh test tubes. A similar protocol was employed when preparing samples for mass spectrometry. Immunoprecipitated proteins were denatured by addition of sample buffer and boiling for 5 minutes, resolved by 8%-16% SDS-PAGE, and analyzed by immunoblotting, CBB staining and silver staining. CBB staining was performed using a Quick-CBB kit (Wako, Osaka, Japan) according to the manufacturer’s instructions. Silver staining was performed using a Sil-Best Stain One kit (Nacalai Tesque, Kyoto, Japan) according to the manufacturer’s instructions.

### Mass spectrometry

Immunoprecipitates from HeLa229, 293T and U2OS cells stably expressing FLAG-PITX1 were prepared as described above. Proteins with the FLAG peptide were eluted from anti-FLAG affinity beads and air dried for nanoLC-MS/MS analysis. Eluted proteins were also detected by CBB or silver staining. Protein pellets were reconstituted in 13 μL of 8 M urea followed by addition of 2 μL of 40 mg/ml DTT. Tubes were gently vortexed and incubated at 37°C for 90 min, and 5 μL of 40 mg/ml IAA was added after reducing, followed by incubation for another 30 min at 37°C in the dark. Then, 60 μl of water was added to tubes. Dilute proteins were digested overnight with 5 μl of 25 ng/μl trypsin gold (Promega, Madison, WI, USA) and desalted with a ZipTipC18 pipette tip (Millipore, Eschborn, Germany), according to the manufacturer’s instructions. Peptides were diluted with water and air dried to remove acetonitrile. Formic acid was added to the peptide solution to a final concentration of 3.3%. The resulting digests were analyzed by DiNa nano liquid chromatography system (KYA Technologies) coupled with a QSTAR Elite hybrid mass spectrometer (AB Sciex, Ontario, Canada). MS/MS spectra were searched against the NCBInr and SwissProt databases using ProteinPilot™ software 2.0 (AB Sciex).

### Transfection and co-immunoprecipitation assay

Cells were transfected with plasmid vectors using Lipofectamine 2000 (Invitrogen). For stable overexpression of FLAG tagged PITX1, 1×10^6^ HeLa229, 293T or U2OS cells were seeded in each well of 6-well plates, and were transfected 24 h after seeding with 0.5 μg of plasmid. Stable cell lines were generated using G418.

Transient transfection was performed with Lipofectamine 2000 according to the manufacturer’s protocol (Invitrogen). Transfected cells were harvested after 24 h in the following ice-cold lysis buffer; 100 mM Tris HCl (pH 7.5), with 300 mM NaCl, 2mM EDTA, 2% Triton X-100, and 400 μl of 25× EDTA-free protease inhibitors (Sigma) per 5 ml. Immunoprecipitation was performed according to the manufacturer’s instructions for the FLAG Immunoprecipitation Kit (Sigma).

### Western blotting analysis

Western blotting was performed as described previously [27]. Membranes were blotted with rabbit polyclonal antibody against human PITX1 antigen (ab70273, 1:2,000; Abcam, Cambridge, MA, UK), rabbit polyclonal antibody against the human ZCCHC10 antigen (HPA038944, 1:2,000; Sigma), mouse monoclonal antibody against the FLAG antigen (F1804, 1:2,000; Sigma), or with polyclonal antibody against β-tubulin (PA1-41331, 1:5,000; Thermo Fisher Scientific, Waltham, MA, USA) and the appropriate standard peroxidase-labeled anti-mouse IgG and anti-rabbit IgG secondary antibody, according to the manufacturer’s instructions (GE Healthcare, Piscataway, NJ, USA). Immunoreactive bands were visualized using the ECL detection system (Thermo Fisher Scientific).

### qRT-PCR

RNA isolation and reverse transcriptase (RT)-PCR was performed as described previously [13]. mRNA expression of *PITX1 hTERT, ZCCHC10* was analyzed using specific primers: *PITX1*: forward; 5’-GCTACCCCGACATGAGCA, reverse; 5’-GTTACGCTCGCGCTTACG, *hTERT*: forward; 5’-GCCTTCAAGAGCCACGTC, reverse; 5’-CCACGAACTGTCGCATGT, *ZCCHC10*: forward; 5’-TGGACTTATGAATGCACAGGAA, reverse; 5’-CTACATTGGTTTCTCCAATGCTT. cDNA was amplified using an Applied Biosystems StepOne thermal cycler system and a SYBR green PCR kit (Applied Biosystems, Foster City, CA, USA). mRNA levels were normalized against *GAPDH* mRNA (PCR primers: forward; 5’-AGCCACATCGCTCAGACAC, reverse; 5’-GCCCAATACGACCAAATCC).

### Tissue samples

Biopsy samples of human melanoma were obtained from the Tottori University Hospital. All materials were obtained with written informed consent, and procedures were approved by the Institutional Review Board of Tottori University (Permission No.1558). All experiments were performed in accordance with the guidelines of the Ethics Committee of Tottori University. Total RNA was extracted from tissue samples using the RNeasy plus kit (Qiagen, Valencia, CA, USA), according to the supplier’s instructions.

### Statistics

Data from more than three separate experiments are presented as means ±S.D. Significance was established at P-values less than 0.05 using an unpaired two-tailed Student’s t-test.

## Acknowledgments

This research was partly performed at the Tottori Bio Frontier managed by Tottori prefecture. We would like to thank Dr. Teruhiko Suzuki for assisting with our FLAG-Pull down examination. This research was partly performed at the Tottori Bio Frontier managed by Tottori prefecture.

Supporting Information Fig S1. Mapping of 57 candidate genes identification of pull-down assay. ‘p’ refers to the short arm of each chromosome, and ‘q’ refers to the long arm. Numbers indicate chromosome number. Red arrow shows 6 genes coding on 5q.

## References

1. Bernardes de Jesus B, Blasco MA. Telomerase at the intersection of cancer and aging. Trends Genet. 2013;29(9):513–20. doi: 10.1016/j.tig.2013.06.007. PubMed PMID: 23876621; PubMed Central PMCID: PMCPMC3896987.

2. Harley CB, Futcher AB, Greider CW. Telomeres shorten during ageing of human fibroblasts. Nature. 1990;345(6274):458–60. doi: 10.1038/345458a0. PubMed PMID: 2342578.

3. Arndt GM, MacKenzie KL. New prospects for targeting telomerase beyond the telomere. Nat Rev Cancer. 2016;16(8):508–24. doi: 10.1038/nrc.2016.55. PubMed PMID: 27339602.

4. Blasco MA. Telomeres and human disease: ageing, cancer and beyond. Nat Rev Genet. 2005;6(8):611–22. doi: 10.1038/nrg1656. PubMed PMID: 16136653.

5. Bodnar AG, Ouellette M, Frolkis M, Holt SE, Chiu CP, Morin GB, et al. Extension of life-span by introduction of telomerase into normal human cells. Science. 1998;279(5349):349–52. PubMed PMID: 9454332.

6. Lee KM, Choi KH, Ouellette MM. Use of exogenous hTERT to immortalize primary human cells. Cytotechnology. 2004;45(1-2):33–8. doi: 10.1007/10.1007/s10616-004-5123-3. PubMed PMID: 19003241; PubMed Central PMCID: PMCPMC3449956.

7. Cong YS, Wright WE, Shay JW. Human telomerase and its regulation. Microbiol Mol Biol Rev. 2002;66(3):407–25, table of contents. PubMed PMID: 12208997; PubMed Central PMCID: PMCPMC120798.

8. Wright WE, Piatyszek MA, Rainey WE, Byrd W, Shay JW. Telomerase activity in human germline and embryonic tissues and cells. Dev Genet. 1996;18(2):173–9. doi: 10.1002/(SICI)1520-6408(1996)18:2<173::AID-DVG10>3.0.CO;2-3. PubMed PMID: 8934879.

9. Daniel M, Peek GW, Tollefsbol TO. Regulation of the human catalytic subunit of telomerase (hTERT). Gene. 2012;498(2):135–46. doi: 10.1016/j.gene.2012.01.095. PubMed PMID: 22381618; PubMed Central PMCID: PMCPMC3312932.

10. Ramlee MK, Wang J, Toh WX, Li S. Transcription Regulation of the Human Telomerase Reverse Transcriptase (hTERT) Gene. Genes (Basel). 2016;7(8). doi: 10.3390/genes7080050. PubMed PMID: 27548225; PubMed Central PMCID: PMCPMC4999838.

11. Kugoh H, Ohira T, Oshimura M. Studies of Tumor Suppressor Genes via Chromosome Engineering. Cancers (Basel). 2015;8(1). doi: 10.3390/cancers8010004. PubMed PMID: 26729168; PubMed Central PMCID: PMCPMC4728451.

12. Qi DL, Ohhira T, Oshimura M, Kugoh H. Human chromosome 5 carries a transcriptional regulator of human telomerase reverse transcriptase (hTERT). Biochem Biophys Res Commun. 2010;398(4):695–701. doi: 10.1016/j.bbrc.2010.07.003. PubMed PMID: 20621064.

13. Qi DL, Ohhira T, Fujisaki C, Inoue T, Ohta T, Osaki M, et al. Identification of PITX1 as a TERT suppressor gene located on human chromosome 5. Mol Cell Biol. 2011;31(8):1624–36. doi: 10.1128/MCB.00470-10. PubMed PMID: 21300782; PubMed Central PMCID: PMCPMC3126332.

14. Lamonerie T, Tremblay JJ, Lanctôt C, Therrien M, Gauthier Y, Drouin J. Ptx1, a bicoid-related homeo box transcription factor involved in transcription of the pro-opiomelanocortin gene. Genes Dev. 1996;10(10):1284–95. PubMed PMID: 8675014.

15. Island ML, Mesplede T, Darracq N, Bandu MT, Christeff N, Djian P, et al. Repression by homeoprotein pitx1 of virus-induced interferon a promoters is mediated by physical interaction and trans repression of IRF3 and IRF7. Mol Cell Biol. 2002;22(20):7120–33. PubMed PMID: 12242290; PubMed Central PMCID: PMCPMC139826.

16. Schubert SW, Kardash E, Khan MA, Cheusova T, Kilian K, Wegner M, et al. Interaction, cooperative promoter modulation, and renal colocalization of GCMa and Pitx2. J Biol Chem. 2004;279(48):50358–65. doi: 10.1074/jbc.M404587200. PubMed PMID: 15385555.

17. Amendt BA, Sutherland LB, Semina EV, Russo AF. The molecular basis of Rieger syndrome. Analysis of Pitx2 homeodomain protein activities. J Biol Chem. 1998;273(32):20066–72. PubMed PMID: 9685346.

18. Furukawa T, Kozak CA, Cepko CL. rax, a novel paired-type homeobox gene, shows expression in the anterior neural fold and developing retina. Proc Natl Acad Sci U S A. 1997;94(7):3088–93. PubMed PMID: 9096350; PubMed Central PMCID: PMCPMC20326.

19. Marcil A, Dumontier E, Chamberland M, Camper SA, Drouin J. Pitx1 and Pitx2 are required for development of hindlimb buds. Development. 2003;130(1):45–55. PubMed PMID: 12441290.

20. Kolfschoten IG, van Leeuwen B, Berns K, Mullenders J, Beijersbergen RL, Bernards R, et al. A genetic screen identifies PITX1 as a suppressor of RAS activity and tumorigenicity. Cell. 2005;121(6):849–58. doi: 10.1016/j.cell.2005.04.017. PubMed PMID: 15960973.

21. Liu DX, Lobie PE. Transcriptional activation of p53 by Pitx1. Cell Death Differ. 2007;14(11):1893–907. doi: 10.1038/sj.cdd.4402209. PubMed PMID: 17762884.

22. Osaki M, Chinen H, Yoshida Y, Ohhira T, Sunamura N, Yamamoto O, et al. Decreased PITX1 gene expression in human cutaneous malignant melanoma and its clinicopathological significance. Eur J Dermatol. 2013;23(3):344–9. doi: 10.1684/ejd.2013.2021. PubMed PMID: 23816528.

23. Nakabayashi M, Osaki M, Kodani I, Okada F, Ryoke K, Oshimura M, et al. PITX1 is a reliable biomarker for predicting prognosis in patients with oral epithelial dysplasia. Oncol Lett. 2014;7(3):750–4. doi: 10.3892/ol.2013.1775. PubMed PMID: 24527083; PubMed Central PMCID: PMCPMC3919858.

24. Lord RV, Brabender J, Wickramasinghe K, DeMeester SR, Holscher A, Schneider PM, et al. Increased CDX2 and decreased PITX1 homeobox gene expression in Barrett’s esophagus and Barrett’s-associated adenocarcinoma. Surgery. 2005;138(5):924–31. doi: 10.1016/j.surg.2005.05.007. PubMed PMID: 16291394.

25. Chen Y, Knösel T, Ye F, Pacyna-Gengelbach M, Deutschmann N, Petersen I. Decreased PITX1 homeobox gene expression in human lung cancer. Lung Cancer. 2007;55(3):287–94. doi: 10.1016/j.lungcan.2006.11.001. PubMed PMID: 17157953.

26. Chen YN, Chen H, Xu Y, Zhang X, Luo Y. Expression of pituitary homeobox 1 gene in human gastric carcinogenesis and its clinicopathological significance. World J Gastroenterol. 2008;14(2):292–7. PubMed PMID: 18186570; PubMed Central PMCID: PMCPMC2675129.

27. Ohira T, Naohiro S, Nakayama Y, Osaki M, Okada F, Oshimura M, et al. miR-19b regulates hTERT mRNA expression through targeting PITX1 mRNA in melanoma cells. Sci Rep. 2015;5:8201. doi: 10.1038/srep08201. PubMed PMID: 25643913; PubMed Central PMCID: PMCPMC4314654.

28. Liu X, Chen Q, Tsai HJ, Wang G, Hong X, Zhou Y, et al. Maternal preconception body mass index and offspring cord blood DNA methylation: exploration of early life origins of disease. Environ Mol Mutagen. 2014;55(3):223–30. doi: 10.1002/em.21827. PubMed PMID: 24243566; PubMed Central PMCID: PMCPMC4547934.

29. Yin SJ, Cho IH, Yang HS, Park YD, Yang JM. Analysis of the peptides detected in atopic dermatitis and various inflammatory diseases patients-derived sera. Int J Biol Macromol. 2018;106:1052–61. Epub 2017/08/24. doi: 10.1016/j.ijbiomac.2017.08.109. PubMed PMID: 28842203.

30. Matthews JM, Sunde M. Zinc fingers--folds for many occasions. IUBMB Life. 2002;54(6):351–5. doi: 10.1080/15216540216035. PubMed PMID: 12665246.

31. Cantor AB, Orkin SH. Coregulation of GATA factors by the Friend of GATA (FOG) family of multitype zinc finger proteins. Semin Cell Dev Biol. 2005;16(1):117–28. Epub 2004/12/15. doi: 10.1016/j.semcdb.2004.10.006. PubMed PMID: 15659346.

32. Izutsu T, Izutsu N, Iwane A, Takada A, Nagasawa T, Kanasugi T, et al. Expression of human telomerase reverse transcriptase and correlation with telomerase activity in placentas with and without intrauterine growth retardation. Acta Obstet Gynecol Scand. 2006;85(1):3–11. PubMed PMID: 16521673.

33. Xi L, Cech TR. Inventory of telomerase components in human cells reveals multiple subpopulations of hTR and hTERT. Nucleic Acids Res. 2014;42(13):8565–77. doi: 10.1093/nar/gku560. PubMed PMID: 24990373; PubMed Central PMCID: PMCPMC4117779.

34. Li M, Chen SM, Chen C, Zhang ZX, Dai MY, Zhang LB, et al. microRNA 299 3p inhibits laryngeal cancer cell growth by targeting human telomerase reverse transcriptase mRNA. Mol Med Rep. 2015;11(6):4645–9. doi: 10.3892/mmr.2015.3287. PubMed PMID: 25634687.

35. Wei G, Luo H, Sun Y, Li J, Tian L, Liu W, et al. Transcriptome profiling of esophageal squamous cell carcinoma reveals a long noncoding RNA acting as a tumor suppressor. Oncotarget. 2015;6(19):17065–80. doi: 10.18632/oncotarget.4185. PubMed PMID: 26158411; PubMed Central PMCID: PMCPMC4627292.

36. Eisenmann KM, Dykema KJ, Matheson SF, Kent NF, DeWard AD, West RA, et al. 5q- myelodysplastic syndromes: chromosome 5q genes direct a tumor-suppression network sensing actin dynamics. Oncogene. 2009;28(39):3429–41. doi: 10.1038/onc.2009.207. PubMed PMID: 19597464.

37. Ebert BL. Genetic deletions in AML and MDS. Best Pract Res Clin Haematol. 2010;23(4):457–61. doi: 10.1016/j.beha.2010.09.006. PubMed PMID: 21130407; PubMed Central PMCID: PMCPMC3032259.

38. Geuens T, Bouhy D, Timmerman V. The hnRNP family: insights into their role in health and disease. Hum Genet. 2016;135(8):851–67. doi: 10.1007/s00439-016-1683-5. PubMed PMID: 27215579; PubMed Central PMCID: PMCPMC4947485.

39. Lu WJ, Lee NP, Kaul SC, Lan F, Poon RT, Wadhwa R, et al. Mortalin-p53 interaction in cancer cells is stress dependent and constitutes a selective target for cancer therapy. Cell Death Differ. 2011;18(6):1046–56. Epub 2011/01/14. doi: 10.1038/cdd.2010.177. PubMed PMID: 21233847; PubMed Central PMCID: PMCPMC3131943.

40. Allan LA, Fried M. p53-dependent apoptosis or growth arrest induced by different forms of radiation in U2OS cells: p21WAF1/CIP1 repression in UV induced apoptosis. Oncogene. 1999;18(39):5403–12. doi: 10.1038/sj.onc.1202931. PubMed PMID: 10498894.

41. Chambers I, Tomlinson SR. The transcriptional foundation of pluripotency. Development. 2009;136(14):2311–22. doi: 10.1242/dev.024398. PubMed PMID: 19542351; PubMed Central PMCID: PMCPMC2729344.

42. Shats I, Milyavsky M, Tang X, Stambolsky P, Erez N, Brosh R, et al. p53-dependent down-regulation of telomerase is mediated by p21waf1. J Biol Chem. 2004;279(49):50976–85. doi: 10.1074/jbc.M402502200. PubMed PMID: 15371422.

43. Kugoh H, Shigenami K, Funaki K, Barrett JC, Oshimura M. Human chromosome 5 carries a putative telomerase repressor gene. Genes Chromosomes Cancer. 2003;36(1):37–47. doi: 10.1002/gcc.10135. PubMed PMID: 12461748.

